# Supplemental LED lighting improves plant growth without affecting biological control in a tri-trophic greenhouse system

**DOI:** 10.1101/2023.04.07.536085

**Authors:** Jessica L. Fraser, Paul K. Abram, Martine Dorais

**Author notes:** Université de Montréal, Institut de recherche en biologie végétale, 4101 rue Sherbrooke Est, Montréal, QC H1X 2B2, Canada. Agriculture and Agri-Food Canada, Saint-Jean-sur-Richelieu Research and Development Centre, 430 Boulevard Gouin, Saint-Jean-sur-Richelieu, QC J3B 3E6, Canada. Corresponding author: Jessica Fraser,.

## Abstract

Artificial lighting, including light-emitting diode (LED) illumination, is increasingly being optimized in protected agricultural systems to maximize plant yield and quality. However, it may also cause other top-down and bottom-up effects in these relatively simple ecological communities that also include insect pests and their natural enemies. While some effects of LED lighting on insects have been demonstrated to date, it is not known how they influence biological control of insect pests in practice. To examine potential top-down and bottom-up impacts of LED illumination on greenhouse biological control with parasitoids, we studied the effects of artificially lengthened days on a tri-trophic system in cages and in a greenhouse. We grew plants under a 12-hour photoperiod of white-supplemented light with 6 hours of additional 1) white light or 2) red and blue light, or 3) with no additional light. We exposed the plants to the pest aphid *Myzus persicae* (Hemiptera : Aphididae) with or without its parasitoid wasp *Aphidius matricariae* (Hymenoptera : Braconidae), or to no insects. The 18-hour light treatments increased mean plant dry mass by 127% compared with the 12-hour control without affecting the aphid’s population density or the parasitoid’s biological control efficacy under relatively low light conditions. This suggests that insect communities in protected agriculture can be resilient to even drastic changes in their light environment, and that adjusting crop lighting in a manner that affects plant growth does not necessarily compromise biological control’s effectiveness.

## Introduction

New technologies have introduced unprecedented environmental conditions into both natural and built environments, including agricultural environments, where they can affect plants, pests, and their natural enemies. One example of this is LED lighting, which provides light with artificial spectra at any time of day. LEDs are used to illuminate fully-indoor plant production systems (Mitchell & Sheibani, 2020) and in greenhouses to provide supplemental light, which is used to compensate for low natural light levels in winter—particularly in northern regions—and maintain yields year-round (Dorais et al., 2017). While high-pressure sodium (HPS) lamps are most commonly used for supplemental lighting in greenhouses, LEDs are emerging as a promising technology because they can provide increased flexibility in use, including their ability to produce highly customizable spectral qualities (Hao et al., 2018; Singh et al., 2015). Given that the spectral quality of crop lighting can influence plant productivity and nutritional quality, and may affect susceptibility to pests, there is incentive to tailor LED growth lighting to produce only the most optimal proportions of each wavelength (Lazzarin et al., 2020; Olle & Viršile, 2013).

Plants, however, are not the only organisms in protected agriculture that are sensitive to the light environment. Pest insects are a major problem in greenhouse production, and biological control is widely used to manage them, with parasitoid Hymenoptera being the largest group of biological control agents used (van Lenteren et al., 2020). Biological control is particularly effective and popular in protected agriculture (Pilkington et al., 2010; Shipp et al., 2007). Both insect herbivores and their natural enemies can respond to artificial spectra, and indeed light is manipulated in pest management, such as for trapping or deterring pests, or disrupting their nocturnal behaviours (Martini et al., 2020; Shimoda & Honda, 2013). In addition to influencing pests, optimizing lighting conditions may allow producers to manipulate control agents’ efficacy (Johansen et al., 2011). For example, providing sufficiently long days can prevent biological control agents from entering diapause (Abrieux et al., 2020). Furthermore, in laboratory or greenhouse settings, some diurnal natural enemies increase their predation and parasitism rates, respectively, under longer photoperiods, or nocturnal ones under shorter photoperiods (Kehoe et al., 2020; Kehoe & van Veen, 2022; Perdikis et al., 1999; Zilahi-Balogh et al., 2006), perhaps because it allows them more time to forage. Some adjust their development time, locomotor activity levels, or offspring sex ratios under particular spectral qualities (Cochard et al., 2017, 2019; Wang et al., 2013).

However, much remains unknown about how LED crop lighting affects greenhouse insects (Johansen et al., 2011). In particular, little is known about how it affects parasitoid wasps (Cochard et al., 2019; Johansen et al., 2011; Lazzarin et al., 2020), despite their ubiquity as greenhouse biological control agents (van Lenteren et al., 2020). To the best of our knowledge, there is no research to date examining how LED crop lighting affects the biological control of pests using parasitoids in a context that also explicitly considers the impacts of the lighting and the biological control on plants as well as insects. Experiments in this vein have been performed to study effects of ecological light pollution (Sanders et al., 2018; Sanders et al., 2015) and effects of shading under natural light (Richards & Coley, 2007; Stoepler & Lill, 2013), which showed that both top-down and bottom-up effects within tri-trophic systems can be influenced by lighting. However, the types of light used in these contexts are very different from the lighting used in protected agriculture. Ultimately, the goal of any biological control program used in managed agroecosystems is to protect plants, so any measures taken to improve biological control must consider not only the effects of lighting on the ability of the biological control agents to suppress pests (i.e. top-down effects), but also how the effects of lighting on plants may exert bottom-up effects on pest populations.

Bottom-up effects can influence agricultural pest populations in various ways (Han et al., 2021), so the effects of supplemental lighting on pests via the host plant should also be considered from the perspective of biological control. Altering the timing or spectral quality of lighting can affect herbivores via changes in plant defenses, nutritional quality, or food availability (Bennie et al., 2015; Limaje et al., 2019; Meijer et al., 2022; Vänninen et al., 2010). In particular, the “plant vigour hypothesis” posits that the fastest-growing plants (or plant parts) will be the most suitable hosts for herbivores (Price, 1991), and phloem-feeding insects tend to be more abundant on them (Cornelissen et al., 2008). Increased sunlight availability can produce increased herbivory in the field (Hough-Goldstein & LaCoss, 2012), so increased daily light integrals from artificial lighting might produce the same effect in greenhouses. Pest densities can also influence their natural enemies’ ability to control them: low densities may limit predators with high food requirements (Osborne et al., 2004), while increases in density can influence the natural enemies’ behaviour according to their functional responses (Fernández-arhex & Corley, 2003). It is possible for top-down effects to override bottom-up effects (Costamagna et al., 2013), but it is not known whether greenhouse LED lighting can produce this result in a parasitoid-host-plant system.

In this experiment we aimed to determine how extending the day with LED lighting of two different spectra impacts the efficacy of greenhouse biological control and plant growth outcomes. We aimed to characterize both top-down effects on aphid populations and plant growth resulting from any changes in parasitoid behaviour, as well as bottom-up effects on pests from the light’s influence on plant growth (Figure 1). As a model system, we used the bell pepper *Capsicum annuum* (Solanales: Solanaceae); the pest aphid *Myzus persicae* (Hemiptera: Aphididae); and the parasitoid *Aphidius matricariae* (Hymenoptera: Braconidae), which is commercially available and used to control *M. persicae* in pepper greenhouses (Blümel, 2004; Boivin et al., 2012). Previously, we showed that *A. matricariae* adjusted its activity timing and intensity in response to the light environment, while keeping total activity levels constant, and showed the most intense activity under light with the highest proportion of short to mid-length wavelengths (i.e. below 600 nm) (Fraser et al., 2023). This is consistent with the fact that most insects—including aphids and most Hymenoptera—have photoreceptors with peak sensitivity to UV, blue, and green light, and would not be expected to see red very well, or far-red at all (Briscoe & Chittka, 2001; Kirchner et al., 2005; Peitsch et al., 1992). In light of this finding, we examined two spectra, both at the same total photon flux density, with one based on a conventional approach to plant production using photosynthetically efficient red and blue LEDs (Cocetta et al., 2017; Massa et al., 2008), and one employing a broader spectrum closer to natural light. We expected that parasitoids would parasitize more aphids and provide better pest control under a broad-spectrum, shorter-wavelength enriched light compared to a red and blue light source, and under LED-extended days compared to non-extended days. Conversely, we expected the plants to perform best (in the absence of aphids) under the red and blue light source, as well as under extended compared to non-extended days. In the absence of parasitoids, the aphids—if the aphids can benefit from bottom-up effects of light on plant growth—were expected to follow the same trend as the plants. We expected to gain insight into how artificial light environments influence plant-herbivore-parasitoid communities, and how LED lighting can impact the ability of parasitoid biological control agents to protect plants in greenhouse plant production.

**Figure 1.**
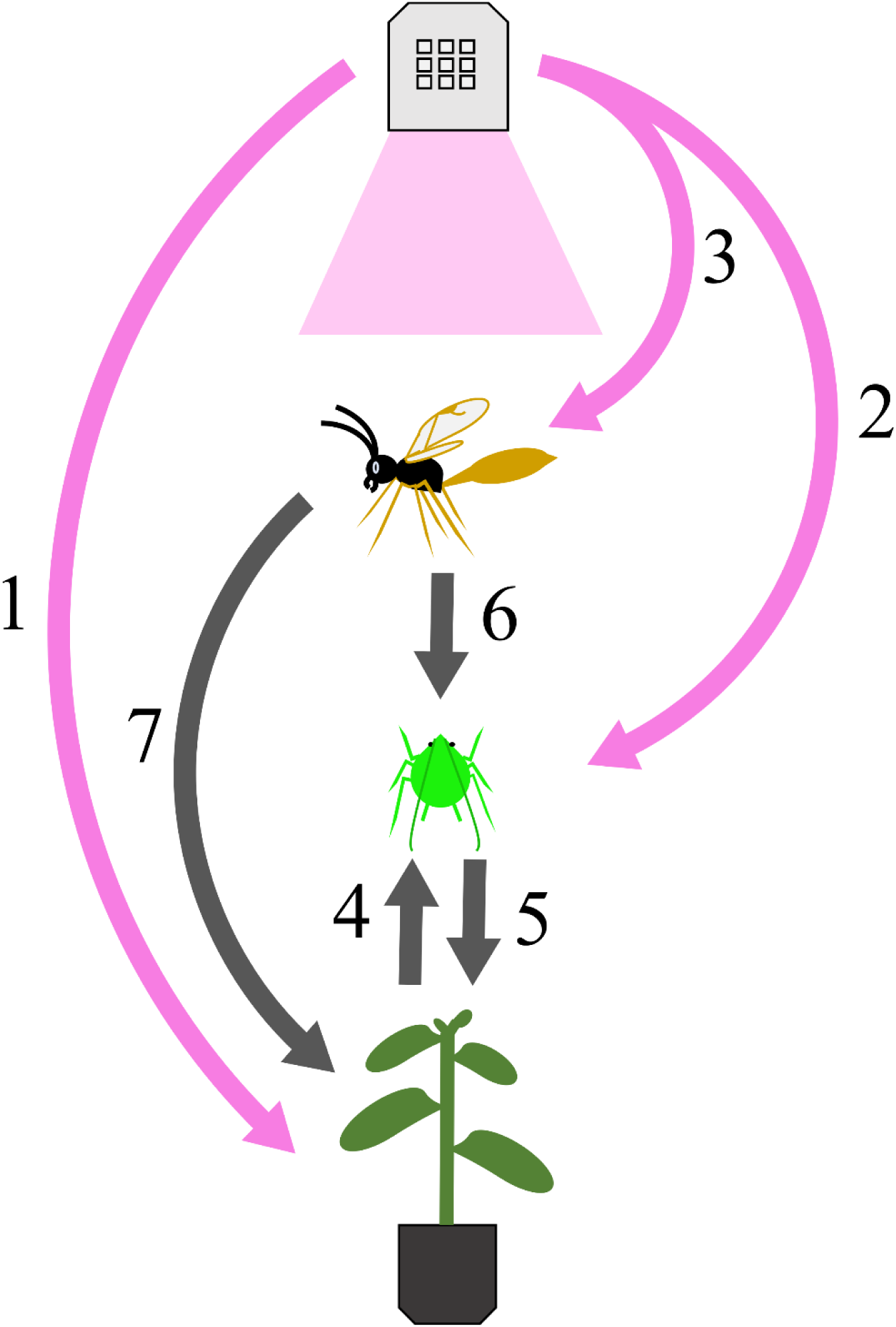
Tri-trophic interactions may be influenced by LED lighting. (1) Increased light availability (photoperiod or intensity) and optimal spectral quality can improve plant growth (Hao et al., 2018). (2) Illumination might have subtle direct effects on aphid fecundity (Joschinski et al., 2015). (3) Choice of lighting may influence parasitoid activity (Cochard et al., 2017; Fraser, 2022; Fraser et al., 2023) and parasitism rates (Cochard et al., 2019; Kehoe et al., 2020). (4) Phloem-feeding insects, like aphids, may benefit from improved plant growth (Cornelissen et al., 2008). (5) Aphids harm plant growth and yield through feeding, as well as transmitting viruses and promoting mold (Dedryver et al., 2010). (6) Parasitoids can control greenhouse aphid populations (Boivin et al., 2012). (7) Parasitoids’ ability to protect greenhouse plants from aphids might be influenced by LED lighting.

## Materials and methods

All experiments were conducted at the Agriculture and Agri-Food Canada Agassiz Research and Development Centre (AAFC ARDC) in Agassiz, British Columbia, Canada (49°14′N, 121°45′W).

### Plant production

*Capsicum annuum* var. Seminis hybrid sweet pepper PS 0941819 (Bayer, Leverkusen, Germany), which used for all experiments and insect rearing, was grown from seed in a peat and perlite medium (pH 5.6– 6.2) and irrigated with an 18N-6P-20K complete fertilizer solution (pH 6.0–6.4) (Poinsettia Plus, Master Plant-Prod, Brampton, Canada). The fertilizer provided 100 ppm N (total), 14 ppm P, 93 ppm K, 11 ppm Mg, 0.560 ppm Fe, 0.280 ppm each Zn and Cu, 0.0560 ppm B, and 0.336 ppm Mo. Plants were kept at target settings of 20–22 ℃ in the daytime and 18–20 ℃ at night, and 50–65% relative humidity. Plants were covered by mesh cages to exclude pests. Sunlight was supplemented with 400–600W HPS lamps (Model LR48877, P.L. Light Systems, Beamsville, Canada; Model LR86176, Light-Tech Systems, Stoney Creek, Canada) at a 16L:8D photoperiod which switched off when natural light levels were above 400 W m^-2^. Light intensity was variable depending on the placement of the plants, but at the height of the canopy, after accounting for absorption by the cages, plants received approximately 50 µmol m^-2^s^-1^ PPFD (average 47.9 µmol m^-2^s^-1^, SE 7.2, based on 10 readings) from the lamps when they were switched on, as measured with an Optimum SRI-2000 handheld spectrophotometer (Optimum OptoElectronics Corp, Hsinchu, Taiwan). The total daily light integral in the pepper growing compartments at the canopy’s height during the course of the experiments was, on average, 2.3 and 2.8 mol m^-2^ d^-1^ respectively, as measured by an Argus Titan OMNI-sensor (version 3.0; Argus Control Systems Ltd., Surrey, Canada) installed in each compartment. These values comprise both sunlight and HPS supplementation, and do not account for light absorption by insect cages.

### Insect rearing

*Myzus persicae* aphids used came from a clonal line maintained at the AAFC ARDC (Uriel et al., 2021) which was maintained on *C. annuum* plants under fluorescent lights (Model F32T8, Philips Lighting, Eindhoven, Netherlands) providing a mean of 40.2 µmol m^-2^ s^-1^ at the height of the canopy, measured with an Optimum SRI-2000 handheld spectrophotometer for a 16L:8D (i.e. 16 hours light, 8 hours dark) photoperiod at target settings of 20 ℃ and 40–60% relative humidity. Aphids used in the experiments were reared outside the main colony to ensure uniformity in age and crowding levels: one green, apterous aphid was placed on a leaf trimmed to fit into a 120 mL polystyrene coffee cup (Loblaw Co. Ltd, Brampton, Canada) with approximately 45 mL of water at the bottom which was covered with a 59 mL plastic SOLO cup (Dart Container Corporation, Toronto, Canada) with a hole in it to allow the leaf’s petiole to pass through. The cup was covered by a plastic coffee cup lid (Dixie, Atlanta, Georgia, USA) with a tissue-covered hole for ventilation. After 6 days, the oldest offspring were used in the experiments.

Parasitoids used were from a colony of *A. matricariae* established from a stock of 500 parasitoids purchased from Biobest (Biobest Group NV, Westerlo, Belgium). Every 4–5 months, we added additional parasitoids purchased from the same source to minimize potential inbreeding depression. We maintained the parasitoids on whole pepper plants infested *M. persicae* from the main colony under fluorescent lights (Model F32T8, Philips Lighting, Eindhoven, Netherlands) providing a mean of µmol 27.4 m^-2^ s^-1^ at the level of the canopy, measured with an Optimum SRI-2000 handheld spectrophotometer for a 16L:8D photoperiod. Target climate settings were 20 ℃ and 40–60% relative humidity. For each new generation of parasitoids, two cages were prepared with three plants each. Eight aphids were gently transferred onto each plant, and after five days, 15 parasitoids were added to the cage, plus honey for food. Nine to eleven days later, newly formed mummies were gently removed from the plant and transferred to a smaller cage with one pepper plant, some droplets of honey, and a cup of water plugged with a cotton wick. These parasitoids were collected on the day they emerged (or on a few occasions of low emergence, the day after) for use in the experiment.

Specimens of both insect species have been preserved at AAFC ARDC to be used as voucher specimens.

### Experimental greenhouse configuration

The goal of our experiment was to assess how LED lighting regimes could affect parasitoid biological control of aphids in a tri-trophic community context, accounting for top-down effects from the light’s influence on parasitoid behaviour and bottom-up effects from the light’s impact on the plants. The experiment was split into two portions: an initial “plant-only” portion where young pepper plants were grown under an LED treatment for three weeks, and an “insect exposure” portion where either aphids, aphids and parasitoids, or no insects were introduced to the plants under the same light treatment for two weeks. This simulated insects colonizing the plants partway through the production cycle.

The experimental greenhouse was divided into nine compartments using opaque curtains which were white on the inside of the compartment to reflect light. In each compartment, we suspended a Heliospectra RX30 programmable LED grow light (Heliospectra Canada Inc., Toronto, Canada) 1.40–1.45 m above the bench top. Sunlight entered through the open top of each compartment, though it was partially shaded by the curtains. Three of the compartments were designated for the plant-only portion of the experiment, and six for the insect-exposure portion. The same compartments were used for each portion throughout, with light treatments randomly assigned within those groupings in each of four temporal blocks, conducted from September 15^th^ to December 21^st^ 2021, with each temporal block lasting five weeks in total. The temporal blocks were staggered so that while the insect-exposure portion of one was underway, the plant-only portion for the next was run simultaneously. For each temporal block, there were two spatial replicates of each LED treatment in the insect-exposure portion, and therefore two replicates of each LED-insect treatment combination, for a total of 8 replicates of each combination across the whole experiment. The temperature in the greenhouse was set to target ranges of 20–22℃ in the daytime and 18–20℃ at night, and the relative humidity was set to 50–65%.

### LED light treatments

The light treatments we used were designed to distinguish how the community responded to the effect of lengthened photoperiod and increased total light, and to the artificial spectral quality used to achieve it. Two spectral qualities were used: a broad-spectrum white (BSW) and a narrowband red and blue (RB) (Figure 2). Spectral distribution plots (Figure 2) were generated using the R package *pavo* (Maia et al., 2019). The BSW treatment was meant as an approximation of natural light and comprised a 5700K white LED supplemented with narrowband UV (peak 380 nm), red (peak 620 nm) and far-red (peak 735 nm) LEDs. The red and blue treatment reflected the fact that red and blue LEDs are widely used in plant production, since red is especially photosynthetically efficient, while blue is important for normal growth responses, and both are particularly energy-efficient (Cocetta et al., 2017; Massa et al., 2008; McCree, 1971). These spectra were configured into three light treatments (Table A-1). All started with 12 hours of illumination by the BSW spectrum, to approximate the median day length at the latitude where the research was conducted (National Research Council Canada, 2020). The lights-on time was set to 05:15, which was before natural sunrise on the longest day to ensure all treatments had the same artificial sunrise. At 17:15, the treatment then transitioned to either 6 additional hours of BSW light (“BSW extension”), 6 hours of RB light (“RB extension”), or no further supplementation (“no extension”). The 12-hour “baseline day” was used to compensate for the reduction in natural light reaching the plants caused by the curtains and cages used to separate the experimental treatments. The total photon flux density of the light treatments was on average about 31 µmol m^-2^ s^-1^ of all photons from 250 to 800 nm at canopy height after accounting for these reductions (Table A-1); this was the highest intensity we could apply while keeping the total photon flux consistent between the two light spectra used. Light measurements were taken with an Optimum SRI 2000 handheld spectrophotometer before sunrise.

**Figure 2.**
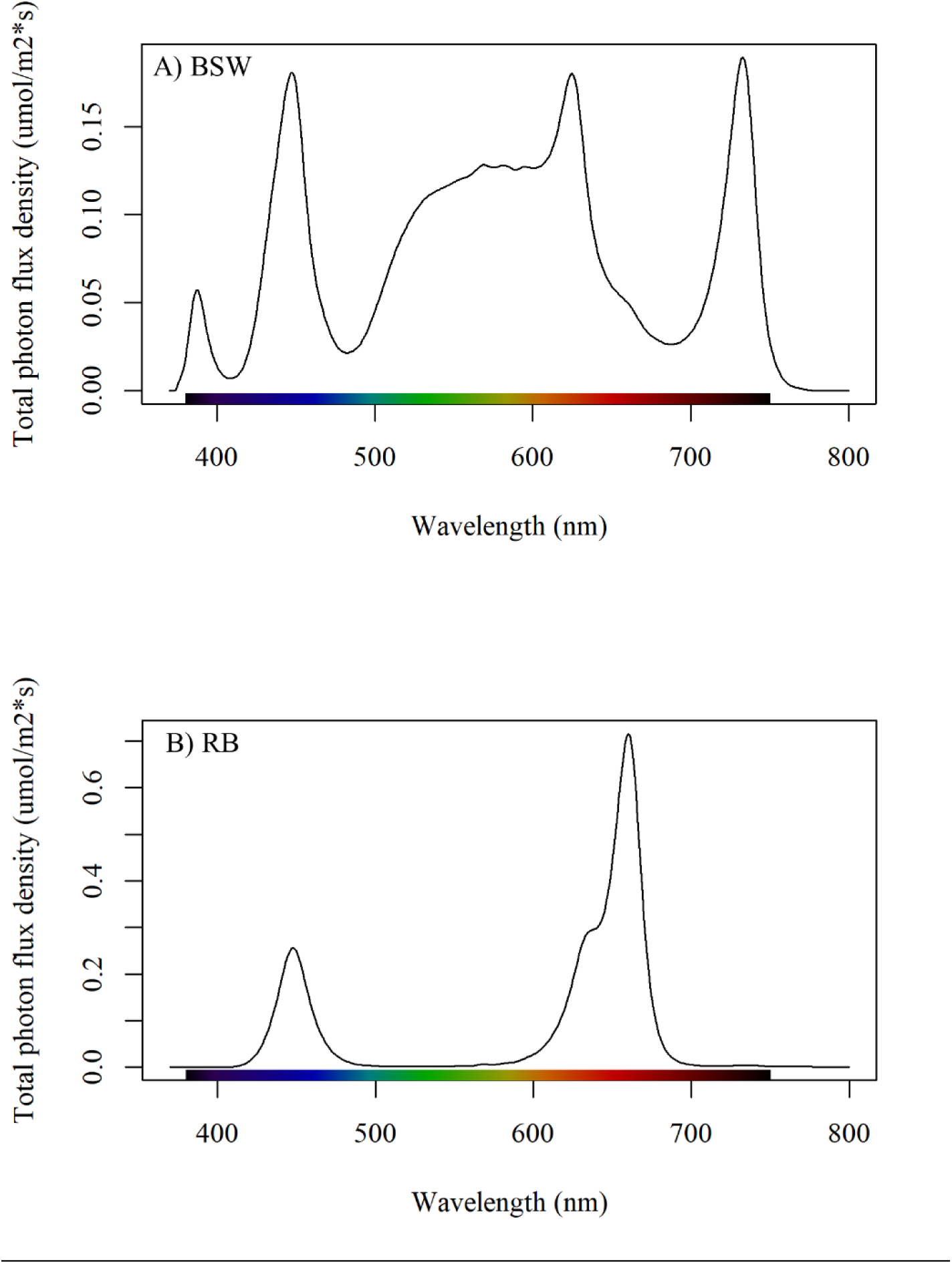
Spectral distributions of light treatments. A) The broad-spectrum white (BSW) setting consisted of 5700K white LED supplemented with UV (380 nm), red (620 nm), and far-red (735 nm) narrowband LEDs. B) The red-blue treatment (RB) comprised blue (450 nm) and red (620 and 660 nm) narrowband LEDs

Artificial light bleeding in from outside the treatment compartments was < 0.75 µmol m^-2^ s^-1^, amounting to 1–2% of the treatment intensity, and came overwhelmingly from HPS lamps in neighbouring greenhouses. Light bleeding into compartments from neighbouring LEDs was below the spectrophotometer’s detection threshold. This HPS spillover occurred during 4.5 of the 6 hours in the “extension” period, when the unextended treatment received no LED light.

Natural light levels decreased as the season progressed. The mean daily light integral from natural photosynthetically active light coming into the greenhouse was approximately 0.8, 0.6, 0.4, and 0.4 mol m^-2^ day^-1^ for the first, second, third, and fourth temporal blocks, respectively, as measured by an Argus Titan OMNI-sensor (version 3.0; Argus Control Systems Ltd., Surrey, Canada) at canopy height, outside the LED compartments. The total daily light integral the plants received, after accounting for absorption by the cages, ranged between 1.4 and 2.1 mol m^-2^ day^-1^ at canopy level depending on the light treatment and natural light levels, as measured with HOBO H22 energy loggers with PAR smart sensors (Onset Computer Corporation, Bourne, United States of America) randomly assigned to compartments during the experiment, placed inside the insect-free control cages at canopy height. Readings taken with the same PAR sensors outside the insect cages but inside LED compartments during the first temporal replicate indicated mean daily light integrals of 2.3–2.4 mol m^-2^ day^-1^. Despite our efforts to compensate for the limited-light conditions, the total daily light integrals in all treatments were substantially lower than the intensities used in commercial pepper production (Dorais et al., 2017).

### Plant growth under LED treatments without insects

Plants were transplanted into 0.5 L pots and transferred into the LED treatments at 16 days post-seeding. Seedlings were randomly assigned to each light treatment and housed in 50 cm x 50 cm x 50 cm mesh insect cages (BugDorm, MegaView Science Ltd., Taiwan), with four seedlings per cage. Drip irrigation lines were fed into the cages and the hole sealed with tape to exclude insects. Plants were grown under these conditions for 19–20 days. Before they were placed in the cages, plant height from the soil top to the shoot apex was measured.

### Plant exposure to aphids and parasitoids under LED treatments

After 19–20 days under their designated LED treatments, the plants were transplanted into 1 L pots and moved to the insect-exposure compartments. LED treatments were randomly assigned to these compartments with two compartments receiving each of the three LED treatments. The plants were randomly assigned to cages within the compartments that had been assigned the LED treatment under which they were initially grown. The plants were housed two to a cage, with the same irrigation tube setup as in the plant-only portion. Within each compartment, three insect treatments were randomly assigned to the cages (one cage per insect treatment): (1) aphids only, (2) aphids and parasitoids, or (3) no insects (control). This produced a factorial split-plot design, with LED treatment as the main plot and insect treatment as the subplot.

Before transfer into the new cages, plant height (soil top to shoot apex) and leaf number were recorded, and the fourth and third true leaves counting from the bottom—or the uppermost full leaves (>2 cm long) if these were not yet developed—were tagged with twist ties loosely fastened around the petiole. The upper one marked where to place the starting aphid, and the lower one marked where to record chlorophyll fluorescence at the end of the experiment. One aphid per plant in each of the insect-treated cages was gently transferred onto the marked starting leaf with a paintbrush.

At two and nine days after the start of the insect exposure portion, five female and five male *A. matricariae*, newly emerged, were added to all of the aphid-parasitoid treatment cages. This was intended to simulate multiple releases into a greenhouse. Only the first group had enough time to produce mummies by the end of the experimental time period, though the second group could still affect them via non-consumptive effects, such as scaring them away from feeding sites (Ingerslew & Finke, 2017). These numbers of parasitoids were used to compensate for the poor survival and low parasitism rates that were observed at lower release rates in preliminary tests.

After 14 (or 15 for the second temporal block) days in the insect exposure portion, for a total of 34 days under the LED treatments, the 49-or 50-day-old plants were removed from the greenhouse and destructively sampled. A two-week insect exposure was chosen to allow the first set of parasitoid offspring to develop to the mummification stage, without allowing aphid populations in the aphid-only cages to become overcrowded.

For the destructive sampling, we took leaf samples for insect density, plant height, leaf number, and above-ground fresh and dry mass for all plants as indicators of growth. We also measured chlorophyll fluorescence F_v_/F_m_ ratio on one randomly selected plant per cage as an indicator of the quantum yield of PSII (Kalaji et al., 2017); this has previously been used in assessing interactions between biotic stress and abiotic stress (Pachu et al., 2021). For the insect sampling, we removed three leaves (>2 cm long) from each plant, comprising the aphid starting leaf, and one each randomly selected above and below it. These were individually removed, weighed, placed into individual bags, and frozen at –18℃. This allowed us to sample aphid and parasitoid mummy numbers throughout the plant canopy (Moerkens et al., 2021; Sanchez et al., 2011). The designated leaf for measuring chlorophyll fluorescence was gently washed with 70% ethanol, then water, and gently dried, to ensure the area was free from aphids and honeydew, and the section to be analyzed was dark adapted for no less than 20 minutes. Chlorophyll fluorescence was then measured using a Hansatech Handy PEA (Hansatech Instruments Ltd., Norfolk, UK) on the predetermined leaf (2^nd^, 3^rd^, or 4^th^ true leaves, counting from the bottom), at the base near the petiole. A saturating light intensity of 3000 µmol m^-2^ s^-2^ was used with a 1 second duration. The plants were then brushed with paper towels to remove any insects still present on the plants. The plant height, remaining leaf number (>2 cm) and above-ground fresh mass (leaf and stem) were recorded. In the third temporal block, the fresh mass, leaf number, and height were recorded the day after experiment takedown for some cages, assuming a negligible time effect. Leaves and stems were dried at 60℃ for no less than 60 hours and dry mass was recorded. The mass of any remaining insects on the leaves was assumed to be negligible. Insect counts were performed on the frozen leaf samples. Numbers of aphids and mummies within each sampling bag were recorded, then leaves were dried and dry mass recorded as for the other samples.

### Statistical analysis

Data were analyzed in R version 4.1.2 (2021-11-01) (R Core Team, 2021). All data were analyzed as the sum of the data collected for both plants within each cage to avoid pseudo-replication, except chlorophyll fluorescence for which only one reading was taken per cage. Some cages were omitted from the analysis: one cage in the BSW-aphid-parasitoid combination, where a plant was accidentally destroyed during transplanting, and four cages in the BSW-aphid-only combination, where parasitoid contamination occurred. A table of sample sizes by treatment is presented in Table A-2. No aphids were observed in any of the control treatment cages.

Using a mixed-effect models to account for the split-plot design with unbalanced data was considered (Crawley, 2013), but the models did not reliably converge, so fixed-effect models were used. All model assumptions were checked by visual inspection of residual plots. Model simplification for all models was performed by assessing the significance of terms within the model through F-tests, and progressively removing non-significant terms, starting with interactions (Crawley, 2013). All models were analyzed with a Type II ANOVA for linear models, or a Type II Analysis of Deviance for the quasi-Poisson and quasi-Binomial models using *car* (Fox & Weisberg, 2019). It was not possible to check for an interaction between temporal block and the other factors in the model (LED treatment and/or insect treatment) due to the low sample size and missing data points. Pairwise comparisons for significant main effects were computed using Tukey’s HSD method with *emmeans* (Lenth, 2022).

Dry mass was log-transformed to meet the assumption of homogeneity of variance. Transformed dry mass was modeled as a function of LED treatment, insect treatment, their interaction, and temporal block. F_v_/F_m_ was analysed with a linear model in the same manner, with no transformation applied. Aphid count data for the aphid-only and aphid-parasitoid treatments (as the sum of all aphids, healthy or mummified) were analyzed with a quasi-Poisson regression model to account for overdispersion (Ver Hoef & Boveng, 2007). The model was expressed as a function of LED treatment, insect treatment, the interaction of LED treatment and insect treatment, and temporal block. The mummification rate in the aphid-parasitoid treatment was analyzed as the proportion of aphids on a leaf that were mummified, as a function of LED treatment and temporal block, using a quasi-binomial regression model to account for overdispersion (Crawley, 2013). The mummification rate underestimated the overall parasitism rate, as it excluded any aphids that were parasitized but had not yet been mummified. However, it should reflect the parasitism rate in the time shortly following the first parasitoid release. For the other plant growth parameters, data were also summed per cage, except chlorophyll fluorescence, for which only one measurement was taken per cage. Height was expressed as the final measured height minus the initial height when plants were moved into the LED treatments at 15–16 days old. The correlation between the total fresh and dry mass, plant height increase, leaf number per cage, and chlorophyll fluorescence (Fv/Fm ratio) were analysed for correlation with a Pearson’s correlogram using *GGally* (Schloerke et al., 2021) (Fig A-2). Fress mass, height increase, and leaf number were strongly correlated with dry mass.

## Results

### Impacts of LED treatments on plant growth

Plant growth was sensitive to the duration and intensity of light available, but not the spectral quality (Figure 3A, Table 1). Fresh and dry mass, height increase, and leaf number were all strongly correlated with one another (correlations of 0.87 to 0.99), but the chlorophyll fluorescence expressed by the F_v_/F_m_ ratio was not substantially correlated with the other measurements (correlations of –0.033 to –0.087) (Fig A-2).

**Figure 3.**
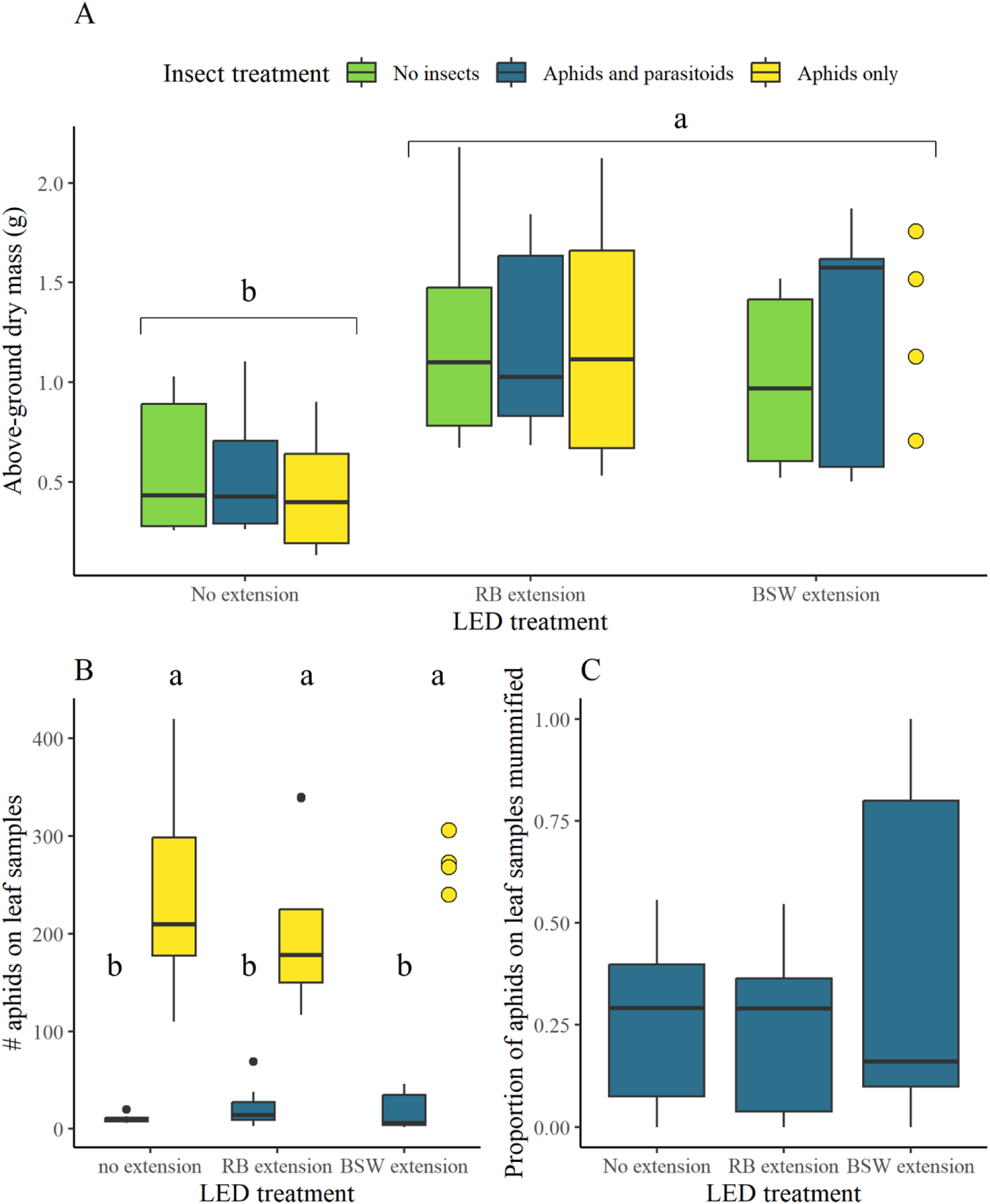
Plant growth, but not aphid population size or parasitism rate, was affected by LED illumination regime. Peppers were grown for five weeks under natural light supplemented with 12-hour days of broad-spectrum white (BSW) light, and then supplemented with another 6 hours of BSW, 6 hours of red and blue (RB) light, or no additional light. For two of those weeks, they were exposed to aphids, aphids and parasitoids, or no insects. (A) Peppers accumulated greater dry mass under both the BSW and RB supplementation treatments, and their dry mass was not affected by aphid infestation. (B) Aphid population size (including mummified and healthy individuals) was reduced by the introduction of parasitoids, but was not affected by the light regime, nor was (C) the proportion of aphids that were mummified by parasitoids. Data are presented as points where n < 5 due to missing data.

**Table 1.** ANOVA results for plant dry mass, aphid population size, and proportion of aphids mummified. Type II ANOVA was performed for the for linear model and Type II Analysis of Deviance for the quasi-family generalized linear models. Rows in italics represent variables removed in model simplification.

| **Response variable** | **Explanatory variable** | **Sum Sq** | **Df** | **F-value** | **p-value** |
| --- | --- | --- | --- | --- | --- |
| Pepper dry mass<br>(log-transformed) | LED treatment | 11.75 | 2 | 142.49 | <0.001 |
|  | Temporal block | 15.51 | 3 | 125.43 | <0.001 |
|  | Residuals | 2.51 | 61 |  |  |
|  | *Insect treatment* | *0.24* | *2* | *3.11* | *0.052* |
|  | *LED treatment : insect treatment* | *0.23* | *4* | *1.54* | *0.20* |
| **Response variable** | **Explanatory variable** | **LR Chisq** | **Df** | **p-value** |  |
| Number of aphids<br>on sampled leaves<br>(quasi-Poisson) | Insect treatment | 195.44 | 1 | <0.001 |  |
|  | *Temporal block* | *6.15* | *3* | *0.10* |  |
|  | *LED treatment* | *0.79* | *2* | *0.67* |  |
|  | *LED treatment : insect treatment* | *2.09* | *2* | *0.35* |  |
| Proportion aphids<br>mummified (quasi-Binomial) | temporal block | 4.08 | 3 | 0.25 |  |
|  | *LED treatment* | *0.85* | *2* | *0.66* |  |

The LED extensions both produced plants with mean dry mass 78% greater than plants grown under non-extended days (Fig 3A, Table 1). The RB treatment produced a slightly (+5%), but not statistically significantly, higher plant dry mass than the BSW (p = 0.052; Table A-4). The plant dry mass decreased substantially in later temporal blocks, in a manner that correlated closely with the reduction in natural light intensity observed over time, amounting to a 61% decrease in mass at the end of the season compared to the start (p < 0.001; Table A-3, Fig A-1). The plant fresh mass, leaf number, and plant height increase were strongly correlated with the plant dry mass (correlation = 0.99, 0.93, 0.92. respectively) and with each other (correlation ≥ 0.87) (Fig A-2), so they increased under conditions of greater light availability. The F_v_/F_m_ ratio was affected by the LED treatment, being higher in the non-extended treatment than the BSW treatment (p < 0.001) and the RB treatment (p < 0.001), though the BSW and RB did not differ from each other (p = 0.069) (Table A-5, Fig A-3). It also differed by temporal block (p < 0.001), though only between the first and third blocks (p = 0.0020) and the second and third blocks (p = 0.0087).

### Impacts of insects on plant growth

We did not detect significant effects of the insects on plant growth. The plant dry mass was not significantly affected by insect treatment (p = 0.052), though the aphid-only treatment had slightly lower dry mass (<= 5% difference) than the control and the parasitoid-exposed treatments. The LED treatment did not modulate the insects’ impact on the plants, either in terms of dry mass (p = 0.20; Table 1), or of F_v_/F_m_ ratio (p = 0.41; Table A-5). Further details are presented in Tables A-4 (dry mass) and A-5 (F_v_/F_m_ ratio).

### Impacts of LED treatments on insect populations

While the plants responded strongly to the light conditions, neither aphid densities (p = 0.67) nor parasitoid mummification rates (p = 0.66) were shown to be affected by the LED treatments (Table 1). However, the presence of parasitoids drastically reduced aphid numbers (p <0.001) regardless of the temporal block start date (p = 0.10) and did not interact with the LED treatment (p = 0.35) (Fig 3B).

Similarly, the proportion of sampled aphids that were mummified was also unaffected by the time within the season (p = 0.25) (Figure 3C). Specific p-values for the mummification rate model should, however, be interpreted cautiously due to the low sample size (Crawley, 2013). Aphid numbers were 93% lower in the parasitoid-exposed cages compared with the aphid-only ones. There were on average 38.6 aphids per leaf in the aphid-only treatment compared to 2.3 non-mummified aphids per leaf and 0.5 mummies per leaf in the parasitoid-exposed treatment. This result suggests that parasitism rates were similar across changes in both artificial lighting treatments and seasonal declines in natural light. Additional details are presented in Table A-3.

## Discussion

Global production of fruits and vegetables will need to increase substantially in the coming decades to satisfy global nutritional requirements, and high-tech, energy-efficient greenhouses and vertical farms that use LED lighting could help to fulfill this need (Kc et al., 2018). It will be crucial to understand how the relatively simple communities in these agroecosystems interact as new technologies emerge that might affect these interactions. Here, we specifically asked how pest suppression (and resultant plant protection) by a biological control agent might be affected by LED lighting, which is increasingly being used in greenhouse production. While photoperiod and spectral quality have previously been shown to affect insect parasitoid behaviour (Cochard et al., 2017, 2019; Fraser et al., 2023; Kehoe et al., 2020; Zilahi-Balogh et al., 2006), it was not known how this translated to impacts on biological control efficacy (i.e. protection of crop plants from insect pests) when artificial LED spectra are used to supplement sunlight in a greenhouse setting. We found that it is possible to strongly alter the light conditions within a greenhouse to improve plant growth without provoking corresponding changes in pest and parasitoid populations.

The LED treatments tested significantly affected plant growth via total light duration under low light conditions, but did not observably impact insect populations. Parasitoids were highly effective in reducing aphid populations regardless of light conditions, and no effects of lighting on aphid numbers in the absence of parasitoids were observed.

### Direct effects of lighting

The LED lighting regimes substantially affected plant growth. The increase we observed in above-ground plant dry mass in the BSW and RB extended days relative to the 12-h control can be ascribed to the increase in daily light integral, rather than the longer photoperiod, similar to the observations of Garcia and Lopez (2020). In other words, LED lighting providing a greater light intensity over a shorter photoperiod would probably have produced the same effect. This is supported by the fact that changes in natural daily light integral over the course of the season produced a similar response, and while these changes affected the intensity of the light the plants received, they did not alter the photoperiod because the LED treatments held it constant. We also found that photosystem II efficiency as measured by chlorophyll fluorescence (F_v_/F_m_) tended to be slightly higher in the non-extended (i.e. lower light) treatment compared to the extended treatments. This has previously been observed in peppers grown in low light (Sui et al., 2012), and may represent low-light acclimation. Overall, growth in the lowest-light conditions (12-hour days and later temporal blocks) was substantially reduced relative to plants receiving more light, as would be expected. The effects of spectral quality on the plants were much less pronounced. The RB treatment was expected to produce a slightly greater plant dry mass than the BSW, since it had most of its light in the optimally photosynthetically efficient red range (McCree, 1971), but the slight difference that we observed was not statistically significant. A clear difference between the two spectra might have been observed at higher LED light intensities or greater durations of the extension treatments allowing the LED treatments to make up a larger proportion of the total daily light integral (Hurt et al., 2019), or under a longer experiment duration.

### Direct effects of lighting on aphids

Despite the plants showing a strong difference in growth between the extended and non-extended days, no effect of LED regime on aphid populations was observed. The same was true for differences in natural light levels among the temporal blocks. We established in previous research (Fraser et al., 2023) that lighting conditions very similar to these LED treatments did not produce significant effects on *M. persicae*’s fecundity. This reinforces our findings (Fraser et al., 2023) that greenhouse LED lighting is not likely to substantially affect the growth of aphid populations once the aphids have settled on a host plant, even in cases where the plants are strongly affected.

### Plant*-insect interactions*

We did not find evidence for plant mass influencing aphid population density, or aphids reducing plant mass under the conditions tested. Aphids, in the absence of parasitoids, did show a marginal tendency to reduce the plant dry mass, hinting that the infestation was severe enough that it may have had a significant effect had the experiment lasted longer. We also did not observe any effect of the aphids on the photosystem II efficiency of the plants as measured by chlorophyll fluorescence (F_v_/F_m_ ratio), which is consistent with the findings of Macedo et al. (2003) in soybean.

We did not see any evidence for changes in plant growth differentially affecting the aphids through bottom-up effects under the LED treatments. There were two ways bottom-up effects from increased light could have manifested: the plants could have provided superior food for aphids under higher-light conditions, in accordance with the plant vigour hypothesis (Hough-Goldstein & LaCoss, 2012), or they could have produced more direct defenses against herbivory. The latter seems unlikely due to the low overall light intensities, which can limit plant investment in defenses (Vänninen et al., 2010). Given the fact that the increase in plants’ growth rate in response to increased daily light integral is the greatest at low light intensities and in younger pepper plants (Bruggink, 1987), if *M. persicae* was particularly sensitive to host plant growth rate, we might have expected to detect it; however, these findings should be re-examined under higher light intensities. Indeed, previous research showed that pea plants grown under low light (0.01 µmol m^-2^ s^-2^) had lower levels of defensive chemicals, lower aphid tolerance, lower levels of photosynthetic pigments, and reduced growth, while aphids feeding on them were shorter-lived and less fecund, compared to plants receiving full fluorescent light (91.99 µmol m^-2^ s^-2^) (Dancewicz et al., 2017). While our low-light conditions were not this extreme, they may nevertheless have masked treatment effects to some extent. It could also be that in a choice scenario, the aphids would be more inclined to choose the more vigorously growing, LED-supplemented plants, since herbivore choice of host plant is an important part of the plant vigour hypothesis (Cornelissen et al., 2008; Price, 1991), leading to different aphid abundances between LED treatments even in the absence of effects on aphid fecundity. However, even if bottom-up effects did increase aphid densities, biological control might counteract it. Pests can end up facing the trade-off of higher attack rates from predators and parasitoids in areas that would otherwise be prime feeding locations (Costamagna et al., 2013; Stoepler and Lill, 2013).

### Effects of lighting on parasitoids

Aphid control by *A. matricariae* appeared to be robust to changes in lighting conditions. We found no differences in its effectiveness at controlling *M. persicae*, either in terms of aphid population reductions or mummification rate, under any of the light treatments studied. This is consistent with our findings in (Fraser et al., 2023) that *A. matricariae* kept its total activity levels constant regardless of the photoperiod or spectral quality. We did see very large reductions in aphid population sizes in parasitoid-exposed cages, as well as some evidence to suggest that the parasitoids had a protective effect on the plant dry mass accumulation. However, our observed mean of 2.3 non-mummified *M. persicae* per leaf in the parasitoid-exposed treatment is within the range detected using a similar leaf-sampling method in unheated commercial pepper greenhouses undergoing IPM (Sanchez et al., 2011), which suggests that the levels of control the parasitoids achieved in our experiment are not unrealistically high.

The observed consistency in parasitoid efficacy across lighting conditions may be species-dependent, since some parasitoids have been found to increase their parasitism under extended photoperiods or dim artificial light at night while others under the same conditions were not (Sanders et al., 2018; Zilahi-Balogh et al., 2006). However, our work offered a much lower aphid-to-parasitoid ratio than previous research, which has tended to provide hosts in large numbers (Cochard et al., 2019; Kehoe & van Veen, 2022; Zilahi-Balogh et al., 2006) or even in excess (Kehoe et al., 2020). In commercial greenhouse production, it is advisable to introduce parasitoids pre-emptively using banker plants, or at least release them as early in the season as possible (Boivin et al., 2012; Payton Miller & Rebek, 2018; Portree & Luczynski, 2004). In this scenario, parasitoids are more likely to contend with smaller numbers of hosts at a time. This suggests that the impact of artificial lighting on parasitoid efficacy may depend on the timing of parasitoid release if its effects tend to become detectable once pest infestations are severe. Overall, our results demonstrated that under relatively low light conditions, *A. matricariae* was capable of controlling *M. persicae* populations regardless of the photoperiod, total daily light integral, or spectral quality that was used to extend the day, indicating that it is possible to make substantial changes in light conditions without compromising biological control effectiveness.

### Conclusions

The continued efficacy of parasitoids under all the light conditions studied, combined with the absence of bottom-up benefits from supplemental lighting to aphids despite clear benefits to the plants, suggests biological control can be resilient to changes in the light environment. While further research is needed for a clearer understanding of the role of spectral quality and longer-term dynamics in a variety of protected agriculture systems, our results reinforce the idea that not all species are strongly affected by disruptions to natural photoperiods and spectral qualities. There are many documented negative impacts of the alteration of the natural light environment on insects by artificial light at night (Desouhant et al., 2019). Our findings support the idea that ecologically speaking, there may be—at least in the short term— “winners” and “losers” to the anthropogenic altering of natural light cycles (Longcore & Rich, 2004) if some species are less severely affected than others. Our findings also imply that sustainable pest management practices such as biological control, even if often considered highly condition-dependent by producers, can remain effective under a variety of light regimes. Assuming our findings hold in other lighting contexts and for other biological control agents, they imply that designers and managers of greenhouse cropping systems can focus on optimizing their lighting regimes for improving plant growth and energy use efficiency, and perhaps promoting beneficial insect attraction or deterring pest colonization of the greenhouse (Martini et al., 2020; Park & Lee, 2021), without having to compromise biological control efficacy.

## Acknowledgements

The authors would like to thank Jason Thiessen for experimental setup and data collection; Clarissa Capko for data collection; Warren Wong for feedback on the experimental design; Yonathan Uriel for insect rearing methods; and Ricardo Sarte, Paige Boegarts, Denika Joiner, and Autumn White for greenhouse support. This research was funded by the Organic Science Cluster 3 on organic farming from Agriculture and Agri-Food Canada in partnership with L’Abri végétal, PremierTech and Inno-3B (grant OSC3-Activity 13) and a Natural Sciences and Engineering Research Council of Canada Graduate Student Master’s scholarship.

## Author contributions

JF, PA, and MD conceived and designed the research. JF and PA conducted experiments. JF and PA analyzed data. JF wrote the manuscript. JF, PA, and MD revised the manuscript. All authors read and approved the manuscript.

## Conflict of interest statement

The authors have no relevant financial or non-financial interests to declare.

## Notes

### Competing Interest Statement

The authors have declared no competing interest.

## References

1. Abrieux, A., Gottlieb, D., & Chiu, J. C. (2020). Manipulation of chronobiology. In D. Ben-Yakir (Ed.), Optical manipulation of arthropod pests and beneficials. CABI.

2. Bennie, J., Davies, T. W., Cruse, D., Inger, R., & Gaston, K. J. (2015, May 5). Cascading effects of artificial light at night: resource-mediated control of herbivores in a grassland ecosystem. Philosophical Transactions of the Royal Society B, 370(1667), 20140131. https://doi.org/10.1098/rstb.2014.0131

3. Blümel, S. (2004). Biological control of aphids on vegetable crops. In K. M. Heinz, R. G. van Dreische, & M. P. Parrella (Eds.), Biocontrol in protected culture (pp. 297-312). Ball Publishing.

4. Boivin, G., Hance, T., & Brodeur, J. (2012). Aphid parasitoids in biological control. Canadian Journal of Plant Science, 92(1), 1–12. https://doi.org/10.4141/cjps2011-045

5. Briscoe, A. D., & Chittka, L. (2001). The evolution of color vision in insects. Annual Review of Entomology, 46, 471–510. https://doi.org/10.1146/annurev.ento.46.1.471

6. Bruggink, G. T. (1987, 1987/05/01/). Influence of light on the growth of young tomato, cucumber and sweet pepper plants in the greenhouse: Calculating the effect of differences in light integral. Scientia Horticulturae, 31(3), 175–183. https://doi.org/10.1016/0304-4238(87)90044-6

7. Cocetta, G., Casciani, D., Bulgari, R., Musante, F., Kołton, A., Rossi, M., & Ferrante, A. (2017). Light use efficiency for vegetables production in protected and indoor environments. The European Physical Journal Plus, 132(1), 43. https://doi.org/10.1140/epjp/i2017-11298-x

8. Cochard, P., Galstian, T., & Cloutier, C. (2017). Light environments differently affect parasitoid wasps and their hosts’ locomotor activity. Journal of Insect Behavior, 30(6), 595–611. https://doi.org/10.1007/s10905-017-9644-y

9. Cochard, P., Galstian, T., & Cloutier, C. (2019, 2019/06/01/). The proportion of blue light affects parasitoid wasp behavior in LED-extended photoperiod in greenhouses: Increased parasitism and offspring sex ratio bias. Biological Control, 133, 9–17. https://doi.org/10.1016/j.biocontrol.2019.03.004

10. Cornelissen, T., Wilson Fernandes, G., & Vasconcellos-Neto, J. (2008). Size does matter: variation in herbivory between and within plants and the plant vigor hypothesis. Oikos, 117(8), 1121–1130. https://doi.org/10.1111/j.0030-1299.2008.16588.x

11. Costamagna, A. C., McCornack, B. P., & Ragsdale, D. W. (2013). Within-plant bottom-up effects mediate non-consumptive impacts of top-down control of soybean aphids. PLOS ONE, 8(2), e56394. https://doi.org/10.1371/journal.pone.0056394

12. Crawley, M. J. (2013). The R Book, Second Edition. Wiley Publishing.

13. Dedryver, C. A., Le Ralec, A., & Fabre, F. (2010, Jun-Jul). The conflicting relationships between aphids and men: a review of aphid damage and control strategies. Comptes Rendus Biologies, 333(6-7), 539–553. https://doi.org/10.1016/j.crvi.2010.03.009

14. Desouhant, E., Gomes, E., Mondy, N., & Amat, I. (2019). Mechanistic, ecological, and evolutionary consequences of artificial light at night for insects: review and prospective. Entomologia Experimentalis et Applicata, 167(1), 37–58. https://doi.org/10.1111/eea.12754

15. Dorais, M., Mitchell, C. A., & Kubota, C. (2017). Lighting greenhouse fruiting vegetables. In R. G. Lopez & E. S. Runkle (Eds.), Light Management in Controlled Environments (pp. 159-169). Meister Media Worldwide.

16. Fernández-arhex, V., & Corley, J. C. (2003). The functional response of parasitoids and its implications for biological control. Biocontrol Science and Technology, 13(4), 403–413. https://doi.org/10.1080/0958315031000104523

17. Fox, J., & Weisberg, S. (2019). An {R} Companion to Applied Regression, Third Edition. Sage. https://socialsciences.mcmaster.ca/jfox/Books/Companion/

18. Fraser, J. L. (2022). Compatibility of LED greenhouse lighting and parasitoid biocontrol for aphid management in pepper [Master’s Thesis, Université Laval]. Québec, Canada. http://hdl.handle.net/20.500.11794/108051

19. Fraser, J. L., Abram, P. K., & Dorais, M. (2023). The effects of LED daylength extensions on the fecundity of the pest aphid Myzus persicae and the daily activity patterns of its parasitoid, Aphidius matricariae. bioRxiv, preprint. https://doi.org/10.1101/2023.03.12.532303

20. Garcia, C., & Lopez, R. G. (2020). Supplemental radiation quality influences cucumber, tomato, and pepper transplant growth and development. HortScience, 55(6), 804–811. https://doi.org/10.21273/hortsci14820-20

21. Han, P., Lavoir, A. V., Rodriguez-Saona, C., & Desneux, N. (2021, Oct 4). Bottom-up forces in agroecosystems and their potential Impact on arthropod pest management. Annual Review of Entomology. https://doi.org/10.1146/annurev-ento-060121-060505

22. Hao, X., Guo, X., Lanoue, J., Zhang, Y., Cao, R., Zheng, J., Little, C., Leonardos, D., Kholsa, S., Grodzinski, B., & Yelton, M. (2018). A review on smart application of supplemental lighting in greenhouse fruiting vegetable production. Acta Horticulturae, 1227. https://doi.org/10.17660/ActaHortic.2018.1227.63

23. Hough-Goldstein, J., & LaCoss, S. J. (2012, 2012/03/01). Interactive effects of light environment and herbivory on growth and productivity of an invasive annual vine, Persicaria perfoliata. Arthropod-Plant Interactions, 6(1), 103–112. https://doi.org/10.1007/s11829-011-9158-z

24. Hurt, A., Lopez, R. G., & Craver, J. K. (2019, 01 Feb. 2019). Supplemental but not photoperiodic lighting increased seedling quality and reduced production time of annual bedding plants. HortScience, 54(2), 289–296. https://doi.org/10.21273/hortsci13664-18

25. Ingerslew, K. S., & Finke, D. L. (2017, Feb 1). Mechanisms underlying the nonconsumptive effects of parasitoid wasps on aphids. Environmental Entomology, 46(1), 75–83. https://doi.org/10.1093/ee/nvw151

26. Johansen, N. S., Vänninen, I., Pinto, D. M., Nissinen, A. I., & Shipp, L. (2011). In the light of new greenhouse technologies: 2. Direct effects of artificial lighting on arthropods and integrated pest management in greenhouse crops. Annals of Applied Biology, 159(1), 1–27. https://doi.org/10.1111/j.1744-7348.2011.00483.x

27. Joschinski, J., Hovestadt, T., & Krauss, J. (2015). Coping with shorter days: do phenology shifts constrain aphid fitness? PeerJ, 3, e1103. https://doi.org/10.7717/peerj.1103

28. Kalaji, H. M., Schansker, G., Brestic, M., Bussotti, F., Calatayud, A., Ferroni, L., Goltsev, V., Guidi, L., Jajoo, A., Li, P., Losciale, P., Mishra, V. K., Misra, A. N., Nebauer, S. G., Pancaldi, S., Penella, C., Pollastrini, M., Suresh, K., Tambussi, E., Yanniccari, M., Zivcak, M., Cetner, M. D., Samborska, I. A., Stirbet, A., Olsovska, K., Kunderlikova, K., Shelonzek, H., Rusinowski, S., & Baba, W. (2017, Apr). Frequently asked questions about chlorophyll fluorescence, the sequel. Photosynthesis Research, 132(1), 13–66. https://doi.org/10.1007/s11120-016-0318-y

29. Kc, K. B., Dias, G. M., Veeramani, A., Swanton, C. J., Fraser, D., Steinke, D., Lee, E., Wittman, H., Farber, J. M., Dunfield, K., McCann, K., Anand, M., Campbell, M., Rooney, N., Raine, N. E., Acker, R. V., Hanner, R., Pascoal, S., Sharif, S., Benton, T. G., & Fraser, E. D. G. (2018). When too much isn’t enough: Does current food production meet global nutritional needs? PLOS ONE, 13(10), e0205683. https://doi.org/10.1371/journal.pone.0205683

30. Kehoe, R., Sanders, D., Cruse, D., Silk, M., Gaston, K. J., Bridle, J. R., & van Veen, F. J. F. (2020). Longer photoperiods through range shifts and artificial light lead to a destabilising increase in host-parasitoid interaction strength. Journal of Animal Ecology, 00, 1–9. https://doi.org/10.1111/1365-2656.13328

31. Kehoe, R., & van Veen, F. J. F. (2022). Longer daylengths associated with poleward range shifts accelerate aphid extinction by parasitoid wasps. Ecological Entomology, 1-6. https://doi.org/10.1111/een.13131

32. Kirchner, S. M., Döring, T. F., & Saucke, H. (2005, Nov). Evidence for trichromacy in the green peach aphid, Myzus persicae (Sulz.) (Hemiptera: Aphididae). Journal of Insect Physiology, 51(11), 1255–1260. https://doi.org/10.1016/j.jinsphys.2005.07.002

33. Lazzarin, M., Meisenburg, M., Meijer, D., van Ieperen, W., Marcelis, L. F. M., Kappers, I. F., van der Krol, A. R., van Loon, J. J. A., & Dicke, M. (2020, 2020/12/22/). LEDs make it resilient: Effects on plant growth and defense. Trends in Plant Science. https://doi.org/10.1016/j.tplants.2020.11.013

34. Lenth, R. V. (2022). emmeans: Estimated Marginal Means, aka Least-Squares Means. In (Version 1.7.2) https://CRAN.R-project.org/package=emmeans

35. Limaje, A., Armstrong, J. S., Paudyal, S., & Hoback, W. (2019). LED grow lights alter sorghum growth and sugarcane aphid (Hemiptera: Aphididae) plant interactions in a controlled environment. Florida Entomologist, 102(1), 174–180. https://doi.org/10.1653/024.102.0128

36. Longcore, T., & Rich, C. (2004). Ecological light pollution. Frontiers in Ecology and the Environment, 2(4), 191–198. https://doi.org/10.1890/1540-9295(2004)002[0191:Elp]2.0.Co;2

37. Maia, R., Gruson, H., Endler, J. A., & White, T. E. (2019). pavo 2: New tools for the spectral and spatial analysis of colour in r. Methods in Ecology and Evolution, 10(7), 1097–1107 https://doi.org/10.1111/2041-210X.13174

38. Martini, X., Funderburk, J. E., & Ben-Yakir, D. (2020). Deterrence of pests. In D. Ben-Yakir (Ed.), Optical manipulation of arthropod pests and beneficials. CABI.

39. Massa, G. D., Kim, H.-H., Wheeler, R. M., & Mitchell, C. A. (2008). Plant productivity in response to LED lighting. HortScience, 43(7), 1951. https://doi.org/10.21273/hortsci.43.7.1951

40. McCree, K. J. (1971). The action spectrum, absorptance and quantum yield of photosynthesis in crop plants. Agricultural Meteorology, 9, 191–216. https://doi.org/10.1016/0002-1571(71)90022-7

41. Meijer, D., Meisenburg, M., van Loon, J. J. A., & Dicke, M. (2022). Effects of low and high red to far-red light ratio on tomato plant morphology and performance of four arthropod herbivores. Scientia Horticulturae, 292. https://doi.org/10.1016/j.scienta.2021.110645

42. Mitchell, C. A., & Sheibani, F. (2020). Chapter 10 - LED advancements for plant-factory artificial lighting. In T. Kozai, G. Niu, & M. Takagaki (Eds.), Plant factory (second edition) (pp. 167-184). Academic Press. https://doi.org/10.1016/B978-0-12-816691-8.00010-8

43. Moerkens, R., Boonen, S., Wackers, F. L., & Pekas, A. (2021, Jun). Aphidophagous hoverflies reduce foxglove aphid infestations and improve seed set and fruit yield in sweet pepper. Pest Management Science, 77(6), 2690–2696. https://doi.org/10.1002/ps.6342

44. National Research Council Canada. (2020). Sunrise/Sunset tables. Retrieved 09-09-2021 from https://nrc.canada.ca/en/research-development/products-services/software-applications/sun-calculator/

45. Olle, M., & Viršile, A. (2013). The effects of light-emitting diode lighting on greenhouse plant growth and quality. Agricultural and Food Science, 22(2), 223–234. https://doi.org/10.23986/afsci.7897

46. Osborne, L. S., Bolckmans, K., Landa, Z., & Peña, J. (2004). Kinds of natural enemies. In K. M. Heinz, R. G. van Dreische, & M. P. Parrella (Eds.), Biocontrol in protected culture (pp. 95-127). Ball Publishing.

47. Pachu, J. K. S., Macedo, F. C. O., da Silva, F. B., Malaquias, J. B., Ramalho, F. S., Oliveira, R. F., & Godoy, W. A. C. (2021, Jan). Imidacloprid-mediated stress on non-Bt and Bt cotton, aphid and ladybug interaction: Approaches based on insect behaviour, fluorescence, dark respiration and plant electrophysiology. Chemosphere, 263, 127561. https://doi.org/10.1016/j.chemosphere.2020.127561

48. Park, Y. G., & Lee, J. H. (2021). UV-LED lights enhance the establishment and biological control efficacy of Nesidiocoris tenuis (Reuter) (Hemiptera: Miridae). PLOS ONE, 16(1), e0245165. https://doi.org/10.1371/journal.pone.0245165

49. Payton Miller, T. L., & Rebek, E.J. (2018). Banker plants for aphid biological control in greenhouses. Journal of Integrated Pest Management, 9(1). https://doi.org/10.1093/jipm/pmy002

50. Peitsch, D., Fietz, A., Hertel, H., de Souza, J., Ventura, D. F., & Menzel, R. (1992). The spectral input systems of hymenopteran insects and their receptor-based colour vision. Journal of Comparative Physiology A, 170, 23–40. https://doi.org/10.1007/BF00190398

51. Perdikis, D. C., Lykouressis, D. P., & Economou, L. P. (1999, 1999/09/01). The influence of temperature, photoperiod and plant type on the predation rate of Macrolophus pygmaeus on Myzus persicae. BioControl, 44(3), 281–289. https://doi.org/10.1023/A:1009959325331

52. Pilkington, L. J., Messelink, G., van Lenteren, J. C., & Le Mottee, K. (2010). “Protected Biological Control” – Biological pest management in the greenhouse industry. Biological Control, 52(3), 216–220. https://doi.org/10.1016/j.biocontrol.2009.05.022

53. Portree, J., & Luczynski, A. (2004). Growing greenhouse peppers in British Columbia: A production guide for commercial growers. Ministry of Agriculture, Food and Fisheries and BC Greenhouse Growers’ Association.

54. Price, P. W. (1991). The plant vigor hypothesis and herbivore attack. Oikos, 62(2), 244–251. https://doi.org/10.2307/3545270

55. R Core Team. (2021). R: A language and environment for statistical computing. In R Foundation for Statistical Computing. https://www.R-project.org/

56. Richards, L. A., & Coley, P. D. (2007). Seasonal and habitat differences affect the impact of food and predation on herbivores: a comparison between gaps and understory of a tropical forest. Oikos, 116(1), 31–40. https://doi.org/10.1111/j.2006.0030-1299.15043.x

57. Sanchez, J. A., La-Spina, M., Michelena, J. M., Lacasa, A., & Hermoso de Mendoza, A. (2011). Ecology of the aphid pests of protected pepper crops and their parasitoids. Biocontrol Science and Technology, 21(2), 171–188. https://doi.org/10.1080/09583157.2010.530641

58. Sanders, D., Kehoe, R., Cruse, D., van Veen, F. J. F., & Gaston, K. J. (2018, Aug 6). Low levels of artificial light at night strengthen top-down control in insect food web. Current Biology, 28(15), 2474–2478 e2473. https://doi.org/10.1016/j.cub.2018.05.078

59. Sanders, D., Kehoe, R., Tiley, K., Bennie, J., Cruse, D., Davies, T. W., van Veen, F. J. F., & Gaston, K. J. (2015, 2015/10/16). Artificial nighttime light changes aphid-parasitoid population dynamics. Scientific Reports, 5(1), 15232. https://doi.org/10.1038/srep15232

60. Schloerke, B., Cook, D., Larmarange, J., Briatte, F., Marbach, M., Thoen, E., Elberg, A., Toomet, O., Crowley, J., Hofmann, H., & Wickham, H. (2021). GGally: Extension to ’ggplot2’. In (Version 2.1.2) https://CRAN.R-project.org/package=GGally

61. Shimoda, M., & Honda, K.-i. (2013, November 01). Insect reactions to light and its applications to pest management. Applied Entomology and Zoology, 48(4), 413–421. https://doi.org/10.1007/s13355-013-0219-x

62. Shipp, L., Elliott, D., Gillespie, D., & Brodeur, J. (2007). From chemical to biological control in Canadian greenhouse crops. In C. Vincent, M. S. Goettel, & G. Lazarovits (Eds.), Biological control: A global perspective (pp. 118-127). CABI. https://doi.org/10.1079/9781845932657.0118

63. Singh, D., Basu, C., Meinhardt-Wollweber, M., & Roth, B. (2015). LEDs for energy efficient greenhouse lighting. Renewable and Sustainable Energy Reviews, 49, 139–147. https://doi.org/10.1016/j.rser.2015.04.117

64. Stoepler, T. M., & Lill, J. T. (2013). Direct and indirect effects of light environment generate ecological trade-offs in herbivore performance and parasitism. Ecology, 94(10), 2299–2310. https://doi.org/10.1890/12-2068.1

65. Sui, X.-l., Mao, S.-l., Wang, L.-h., Zhang, B.-x., & Zhang, Z.-x. (2012). Effect of low light on the characteristics of photosynthesis and chlorophyll a fluorescence during leaf development of sweet pepper. Journal of Integrative Agriculture, 11(10), 1633–1643. https://doi.org/10.1016/s2095-3119(12)60166-x

66. Uriel, Y., Abram, P. K., & Gries, G. (2021). Parasitoid pressure does not elicit defensive polyphenism in the green peach aphid. Ecological Entomology, 46(3), 668–676. https://doi.org/10.1111/een.13014

67. van Lenteren, J. C., Alomar, O., Ravensberg, W. J., & Urbaneja, A. (2020). Biological control agents for control of pests in greenhouses. In M. L. Gullino, R. Albajes, & P. C. Nicot (Eds.), Integrated pest and disease management in greenhouse crops (pp. 409-439). Springer International Publishing. https://doi.org/10.1007/978-3-030-22304-5_14

68. Vänninen, I., Pinto, D. M., Nissinen, A. I., Johansen, N. S., & Shipp, L. (2010). In the light of new greenhouse technologies: 1. Plant-mediated effects of artificial lighting on arthropods and tritrophic interactions. Annals of Applied Biology, 157(3), 393–414. https://doi.org/10.1111/j.1744-7348.2010.00438.x

69. Ver Hoef, J. M., & Boveng, P. L. (2007). Quasi-Poisson vs. negative binomial regression: How should we model overdispersed count data? Ecology, 88(11), 2766–2772. https://doi.org/10.1890/07-0043.1

70. Wang, S., Tan, X. L., Michaud, J. P., Zhang, F., & Guo, X. (2013, 2013/10/01). Light intensity and wavelength influence development, reproduction and locomotor activity in the predatory flower bug Orius sauteri (Poppius) (Hemiptera: Anthocoridae). BioControl, 58(5), 667–674. https://doi.org/10.1007/s10526-013-9522-2

71. Zilahi-Balogh, M. G., Shipp, J. L., Cloutier, C., & Brodeur, J. (2006). Influence of light intensity, photoperiod, and temperature on the efficacy of two aphelinid parasitoids of the greenhouse whitefly. Environmental Entomology, 35(3), 581–589. https://doi.org/10.1603/0046-225X-35.3.581

