## Supplementary Information: Tables A-1 to A-5, Figures A-1 to A-3 for "Supplemental LED lighting improves plant growth without affecting biological control in a tri-trophic greenhouse system"

**Table A-1 Spectral composition, photosynthetic photon flux density (PPFD), and total photosynthetic photon flux density (TPFD) of light treatments used in the greenhouse experiment.** The greenhouse compartments ("C") were numbered 1–9, and light treatments were assigned randomly to compartments 1–6 (insect exposure portion) and 7–9 (plant-only portion) for each of four temporal replicates. The broad-spectrum white ("BSW") treatment was used in all compartments as the 12-hour baseline day, and in the BSW extension treatment. The red-blue ("RB") treatment was used in the RB extension treatment compartments. Data presented are the mean of three readings in each compartment, one in the position of each of the three cages placed inside the compartment. Light readings were taken with an Optimum SRI-2000 handheld spectrophotometer (Optimum OptoElectronics Corp, Hsinchu, Taiwan). The spectrophotometer was placed 31 cm above the benchtop, which is approximately canopy height, and the readings were taken inside the mesh cages used in the experiment (BugDorm, MegaView Science Ltd., Taiwan) before sunrise.

| | **Mean photon flux density (μmol $m^{-2}$ $s^{-1}$) by wavelength range** | | | | | | |
|---|---|---|---|---|---|---|---|
| **Spectrum and compartment** | **UV (<400 nm)** | **Blue (400-499.5 nm)** | **Green (500-599.5 nm)** | **Red (600-699.5 nm)** | **Far-red (>=700 nm)** | **PPFD (400-700 nm)** | **TPFD (250-800 nm)** |
| **BSW** | **0.8** | **6.2** | **10.9** | **8.2** | **5.2** | **25.3** | **31.2** |
| C1 | 0.7 | 5.8 | 10.6 | 7.5 | 5.0 | 23.9 | 29.6 |
| C2 | 0.7 | 6.6 | 11.5 | 8.3 | 5.0 | 26.3 | 32.0 |
| C3 | 0.8 | 6.2 | 10.3 | 9.1 | 5.1 | 25.5 | 31.4 |
| C4 | 0.9 | 6.2 | 10.8 | 8.3 | 5.3 | 25.3 | 31.5 |
| C5 | 0.8 | 5.9 | 10.9 | 8.0 | 5.3 | 24.9 | 31.0 |
| C6 | 0.8 | 6.0 | 10.7 | 8.0 | 5.0 | 24.7 | 30.4 |
| C7 | 0.8 | 6.1 | 11.1 | 8.0 | 5.3 | 25.2 | 31.3 |
| C8 | 0.8 | 6.2 | 11.2 | 8.1 | 5.1 | 25.5 | 31.4 |
| C9 | 0.8 | 6.4 | 11.4 | 8.5 | 5.2 | 26.2 | 32.3 |
| **RB** | **0.0** | **6.9** | **0.5** | **23.2** | **0.2** | **30.6** | **30.8** |

| | | | | | | | |
|---|---|---|---|---|---|---|---|
| C1 | 0.0 | 7.1 | 0.2 | 22.4 | 0.1 | 29.7 | 29.8 |
| C2 | 0.0 | 6.7 | 0.3 | 24.6 | 0.2 | 31.6 | 31.8 |
| C3 | 0.1 | 7.0 | 1.0 | 22.9 | 0.4 | 30.9 | 31.4 |
| C4 | 0.0 | 6.1 | 0.4 | 23.7 | 0.1 | 30.2 | 30.3 |
| C5 | 0.0 | 7.4 | 0.4 | 22.9 | 0.1 | 30.7 | 30.8 |
| C6 | 0.0 | 6.5 | 0.4 | 22.9 | 0.1 | 29.7 | 29.8 |
| C7 | 0.0 | 7.7 | 0.4 | 23.9 | 0.1 | 32.0 | 32.1 |
| C8 | 0.0 | 6.7 | 1.0 | 20.2 | 0.4 | 27.8 | 28.3 |
| C9 | 0.0 | 7.0 | 0.4 | 25.3 | 0.2 | 32.7 | 32.9 |

**Table A-2 Total sample sizes by treatment combination for greenhouse experiments.** Four cages in the BSW-aphid-only treatment combination were lost to parasitoid contamination: one in the second temporal block, one in the third, and two (both) in the fourth. One cage in the BSW-aphid-parasitoid treatment combination was lost due to the accidental destruction of a plant in the fourth temporal block.

| Treatment | BSW extension | RB extension | No extension |
|---|---|---|---|
| Control (no insects) | 8 | 8 | 8 |
| Aphid-only | 4 | 8 | 8 |
| Aphid-parasitoid | 7 | 8 | 8 |

**Table A-3 Summaries of greenhouse experiment models.** Estimates and standard deviations obtained by fitting generalized linear models of *Capsicum annuum* dry biomass, *Myzus persicae* density, or proportion of *M. persicae* mummified by *Aphidius matricariae* are displayed. Models were specified as response variable ~ LED treatment*insect treatment + temporal block. Italicized rows indicate factors that were removed in stepwise model selection. $R^2$ and deviance values indicated are for the final model

| Response variable | Parameter | Estimate of fixed effect | Standard error |
|---|---|---|---|
| Plant total dry mass (g) (log-transformed) | Intercept | 0.49 | 0.060 |
| | LED treatment = no extension | -0.78 | 0.063 |
| | LED treatment = RB | 0.15 | 0.063 |
| | Temporal block = 2 | -0.15 | 0.069 |
| | Temporal block = 3 | -1.03 | 0.069 |
| | Temporal block = 4 | -1.04 | 0.071 |
| | *Insect treatment = control* | *0.13* | *0.060* |
| | *Insect treatment = parasitoid* | *0.14* | *0.060* |
| | *LED treatment = no extension : Insect treatment = control* | *0.33* | *0.15* |
| | *LED treatment = RB : Insect treatment = control* | *0.068* | *0.15* |
| | *LED treatment = no extension : Insect treatment = parasitoid* | *0.23* | *0.16* |
| | *LED treatment = RB : Insect treatment = parasitoid* | *0.018* | *0.16* |
| | Multiple $R^2$ = 0.92 | | Adjusted $R^2$ = 0.91 |
| Aphid count (quasi-Poisson) | Intercept | 5.45 | 0.073 |
| | Insect treatment = parasitoid | -2.61 | 0.26 |
| | *Temporal block = 2* | *-0.13* | *0.18* |
| | *Temporal block = 3* | *-0.26* | *0.19* |
| | *Temporal block = 4* | *-0.50* | *0.22* |
| | *LED treatment = no extension* | *-0.043* | *0.19* |
| | *LED treatment = RB* | *-0.16* | *0.19* |
| | *LED treatment = no extension : Insect treatment = parasitoid* | *-0.55* | *0.70* |
| | *LED treatment = RB : Insect treatment = parasitoid* | *0.38* | *0.59* |
| | *Null deviance: 5775.24 on 42 df* | | *Residual deviance: 940.52 on 41 df* |
| Proportion aphids mummified | Intercept | -1.96 | 0.55 |
| | Temporal block =2 | 0.77 | 0.90 |
| | Temporal block = 3 | 0.068 | 0.77 |

| | | | |
|---|---|---|---|
| (quasi-binomial) | Temporal block = 4 | 1.15 | 0.68 |
| | *LED treatment = no extension* | *-0.04* | *0.85* |
| | *LED treatment = RB* | *-0.59* | *0.81* |
| | Null deviance: 86.075 on 22 df | Residual deviance:70.123 on 19 df | |

**Table A-4 Results of pairwise comparisons in effects of factors influencing plant above-ground dry biomass.** Tests were performed using Tukey's HSD. For each response variable, values were averaged across the values of the other main effects. Results are presented on the transformed (i.e. log) scale

| Explanatory variable | Comparison | Estimate | SE | df | t-ratio | p-value |
|---|---|---|---|---|---|---|
| LED treatment | BSW – no extension | 0.78 | 0.063 | 61 | 12.47 | <.0001 |
| | BSW – RB | -0.15 | 0.063 | 61 | -2.40 | 0.051 |
| | no extension – RB | -0.93 | 0.059 | 61 | -15.89 | <.0001 |
| Temporal replicate | 1 – 2 | 0.15 | 0.07 | 61 | 2.20 | 0.13 |
| | 1 – 3 | 1.03 | 0.07 | 61 | 14.97 | <.0001 |
| | 1 – 4 | 1.04 | 0.07 | 61 | 14.55 | <.0001 |
| | 2 – 3 | 0.88 | 0.07 | 61 | 12.60 | <.0001 |
| | 2 – 4 | 0.89 | 0.07 | 61 | 12.31 | <.0001 |
| | 3 – 4 | 0.01 | 0.07 | 61 | 0.14 | 1.00 |

**Table A-5 ANOVA and pairwise comparison results for chlorophyll *a* fluorescence $F_v/F_m$ ratio.** Type II ANOVA was performed on the scaled data. Italicized rows indicate factors that were removed in stepwise model selection. Pairwise comparisons were performed using Tukey's HSD

| **Response variable** | **Explanatory variable** | **Sum sq** | **Df** | **F-value** | **p-value** |
|---|---|---|---|---|---|
| $F_v/F_m$ | LED treatment | 0.00091 | 2 | 24.34 | <0.001 |
| | Temporal replicate | 0.00034 | 3 | 6.00 | 0.001 |
| | Residuals | 0.0011 | 61 | | |
| | *Insect treatment* | *0.000034* | *2* | *0.90* | *0.41* |
| | *LED treatment:Insect treatment* | *0.00016* | *4* | *2.34* | *0.066* |

| **Explanatory variable** | **Comparison** | **Estimate** | **SE** | **df** | **t-ratio** | **p-value** |
|---|---|---|---|---|---|---|
| LED treatment | BSW – no extension | -0.0090 | 0.0013 | 61 | -6.73 | <0.001 |
| | BSW – RB | -0.0030 | 0.0013 | 61 | -2.26 | 0.069 |
| | No extension – RB | 0.0060 | 0.0013 | 61 | 4.78 | <0.001 |
| Temporal block | 1 – 2 | 0.00065 | 0.0015 | 61 | 0.44 | 0.97 |
| | 1 – 3 | 0.0055 | 0.0015 | 61 | 3.78 | 0.0020 |
| | 1 – 4 | 0.0034 | 0.0015 | 61 | 2.24 | 0.13 |
| | 2 – 3 | 0.0049 | 0.0015 | 61 | 3.29 | 0.0087 |
| | 2 – 4 | 0.0028 | 0.0015 | 61 | 1.79 | 0.29 |
| | 3 – 4 | -0.0021 | 0.0015 | 61 | -1.39 | 0.51 |

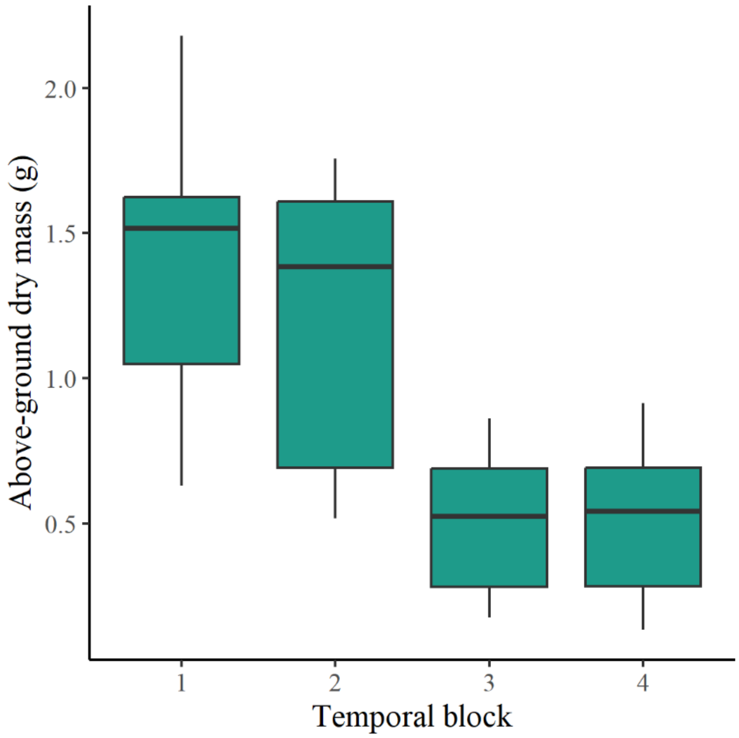


**Figure A-1 Above-ground pepper dry mass per cage by temporal replicate in the greenhouse experiment.** Plants were grown under sunlight supplemented with one of three LED treatments for five weeks, and were exposed to one of three insect treatments. Natural light levels declined as the season progressed. Block start dates and mean daily light integral from sunlight entering the greenhouse over the 5-week block duration were (1) 15 September, 0.8 mol $m^{-2}$ $d^{-1}$; (2) 6 October, 0.6 mol $m^{-2}$ $d^{-1}$; (3) 27 October, 0.4 mol $m^{-2}$ $d^{-1}$ and (4) 17 November, 0.4 mol mol $m^{-2}$ $d^{-1}$ at canopy level (Table B-2), as measured by an Argus Titan OMNI-sensor (version 3.0; Argus Control Systems Ltd., Surrey, Canada) These daily light integral values do not account for light absorption by the insect cages that were used to contain the plants

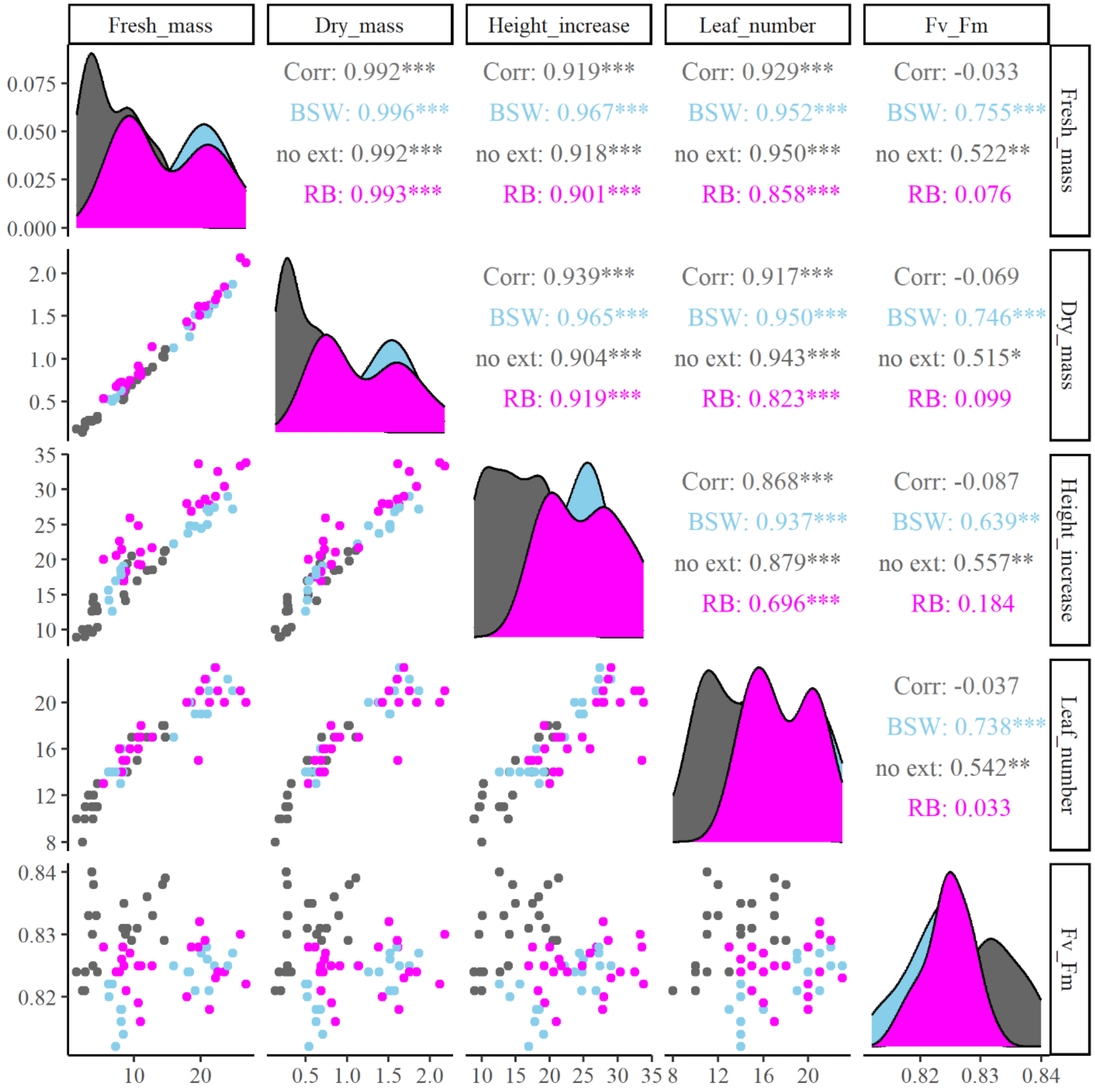


**Figure A-2 Correlation of plant growth parameters by LED treatment in greenhouse experiment.** Plants were grown under sunlight supplemented with one of three LED treatments (broad-spectrum white extension, “BSW”: red-blue extension, “RB”; or no extension, “no ext”) for five weeks, and were exposed to one of three insect treatments. Data presented are the sum of the data collected per cage for dry mass, fresh mass, height increase, and leaf number, except for the $F_v/F_m$ ratio, which was only measured on one plant per cage

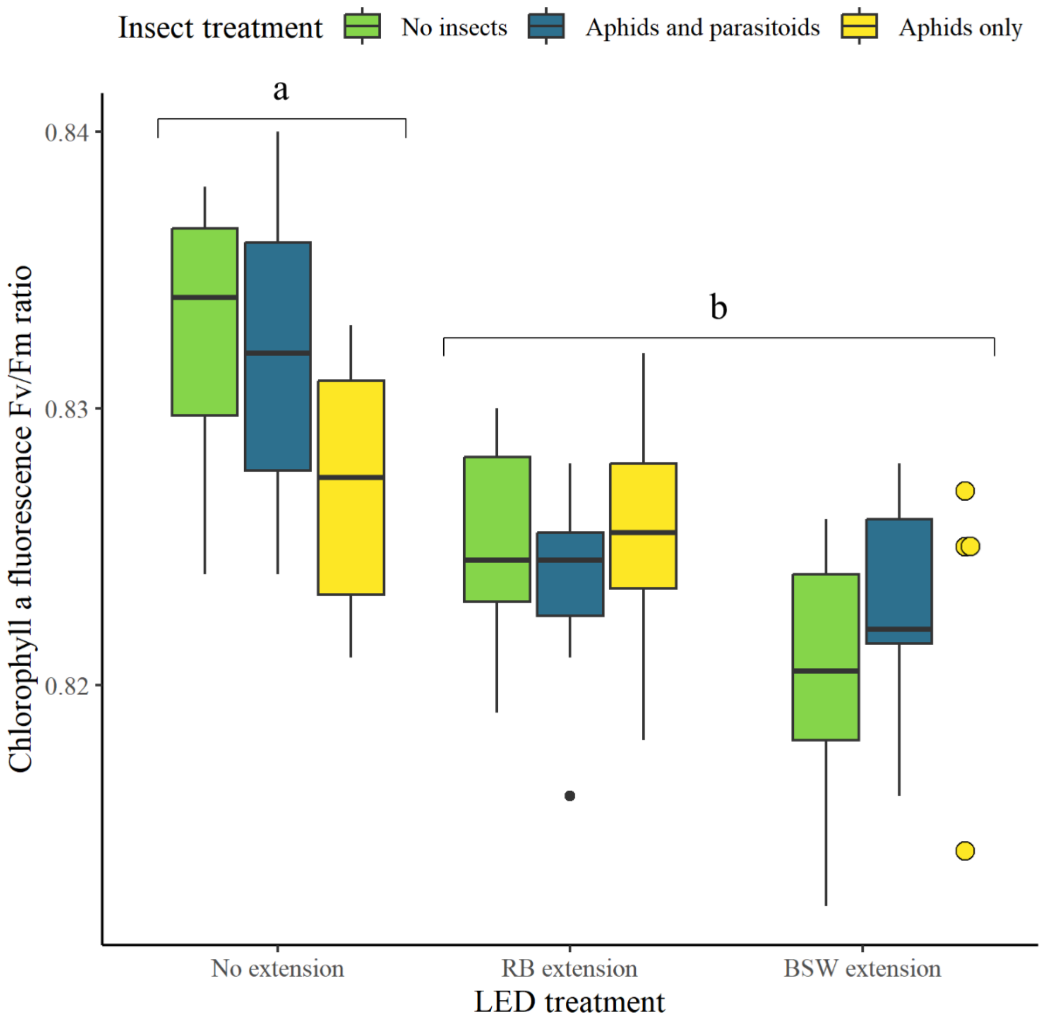


**Figure A-3 Chlorophyll *a* fluorescence $F_v/F_m$ ratio in the greenhouse experiment.** Plants were grown under sunlight supplemented with one of three LED treatments: a broad-spectrum white 12 h day ("No ext"), a broad-spectrum white 18 h day ("BSW extension), or a broad-spectrum white 12 h day plus six hours of red-blue illumination (RB) for five weeks. Plants were exposed to *Myzus persicae* ("Aphids only"), *M. persicae* and *Aphidius matricariae* ("Aphids and parasitoids"), or no insects as a control ("No insects") for two weeks. Significance letters displayed are for Tukey HSD pairwise tests. Data are presented as points where n < 5
